# The data-index: an author-level metric that values impactful data and incentivises data sharing

**DOI:** 10.1101/2020.10.20.344226

**Authors:** Amelia S C Hood, William J Sutherland

## Abstract

Author-level metrics are a widely used measure of scientific success. The h-index, and its variants, measure publication output (number of publications) and impact (number of citations), and these are often used to allocate funding or jobs. Here we argue that the emphasis on publication output and impact hinders progress in the fields of ecology and evolution as it disincentivises two fundamental practices: generating long-term datasets and sharing data. We describe a new author-level metric, the data-index, which values dataset output and impact and promotes generating and sharing data as a result. It is designed to complement other metrics of scientific success, as scientific contributions are diverse and our value system should reflect that. Future work should focus on designing alternative metrics that value our wider merits, such as communicating our research, informing policy, mentoring other scientists, and providing open-access code and tools.

## Introduction

Measuring scientific success with author-level metrics has, despite many concerns, become widely used, including in funding and job allocations (Mingers & Leydesdorff, 2015). The h-index is the most common means of comparison; it combines the publication output (number of publications) and impact (as number of citations) of each scientist (Hirsch, 2005). This metric, despite its many flaws, is a simple and easily calculable measure of scientific impact, so has flourished. Scientists have designed similar metrics that address some of its flaws (Gasparyan et al., 2018; Mingers & Leydesdorff, 2015), such as encouraging unwarranted self-citations (Senanayake, Piraveenan, & Zomaya, 2015), but these metrics are not yet as widely adopted as the h-index. Such flaws are not the focus of this paper. Here, we argue that any value system that solely focuses on publication output and impact will hinder scientific progress.

Though simplicity is alluring, we need a value system that captures our wider merits, such as communicating our research, informing policy, mentoring other scientists, providing open-access code and tools, and generating useful datasets with long-term value. Some of these wider merits are starting to be quantified, for example the field of Altmetrics quantifies the online impact of scholarly work, which is one aspect of science communication (Sud & Thelwall, 2014), and Barres (2013) suggested a metric for measuring mentoring quality. However, many aspects of scientific success are undervalued and if then disincentivised this may slow the progress and impact of science.

Here, we present a new metric, the data-index, designed to value dataset output and impact and complement other metrics of scientific success, such as publication output and impact. Ignoring this dimension of scientific success is particularly detrimental in fields that require expensive, arduous, long-term experiments, such as ecology and evolutionary biology (Hughes et al., 2017; Mills et al., 2015). This is because generating and sharing such datasets is undervalued and even disincentivized. Nearly 50% of scientists surveyed by Kuebbing et al. (2018) cited systemic barriers within academia as hindering efforts to conduct long-term experiments, with pressure to publish frequently as a major cause. This demonstrates the negative impact of our current value system for generating long-term datasets, which contribute disproportionately both scientific understanding and to policy (Hughes et al., 2017).

In terms of data sharing, we first explain the issue from the context of collectors of original data (data generators) sharing their data with data synthesists. Biologists may spend years collecting long-term datasets, only for data synthesists to gain most of the citation credit by reanalysing their work. In the existing system (the h-index), data generators collect “points” for their own publications, but often not for their contribution to syntheses as they are rarely listed as co-authors. Synthesists contend that the citation that they give the generator’s paper is sufficient credit, that generating the data does not meet authorship requirements, or that listing all generators as co-authors would be infeasible. This is often not enough to placate the generators, who may also have concerns that their publication with be overlooked once a synthesis paper becomes available. This hinders scientific progress, as it disincentivises data sharing. In fact, synthesists rarely have success in retrieving data when requested, with an estimated success rate of 50-75% if the researcher is known personally, but lower if not (Côté, Curtis, Rothstein, & Stewart, 2013). Our current system causes authorship disputes that result in important data being left out of synthesis publications. As data synthesists ourselves, we agree that synthesis papers should be cited if available, and citing all of the papers within them is usually infeasible. Therefore, we need a new approach.

Valuing data sharing is important in the context of many types of research, not just synthesis, as it is also important for researchers that want to conduct reanalysis or incorporate other’s datasets into their own. Furthermore, it is essential for achieving scientific reproducibility and it also saves costs by avoiding the unnecessary duplication of results (Piwowar, Vison, & Whitlock, 2011). In light of this, many journals now have policies that mandate public data archiving, but the majority of authors in ecology and evolution (estimated 64% by Roche, Kruuk, Lanfear, & Binning (2015)) archive their data in a way that prevents reuse. This practice is probably partly due to concern over being properly acknowledged when the data is reused, and this is enough to prevent 63% of principle investigators (PIs) from submitting long-term studies to journals that required data archiving (Mills et al., 2015). This demonstrates the need for an improved value system that better rewards collaboration and encourages the archiving of data in a useful manner.

## The Data-Index

The data-index is calculated the same as the h-index is, but it ranks original datasets in order of their citations rather than ranking publications (fig. 1). Dataset citations can come from citations of the original paper that contains the data, or from citations of papers that have numerically re-evaluated the data in the original paper. Figure 1 shows a hypothetical example of how a data generator and synthesist at a similar stage might differ; the generator has a lower h-index than the synthesist, as their publications have fewer citations, but their data-index is higher because they have more original datasets, many of which have been used in several publications. As with any metric of scientific success there will still be some biases, for example when comparing scientists in different fields (Kokko & Sutherland, 1999). However, this worked example demonstrates how using multiple complementary indices in tandem can create a more balanced view of the diversity of scientific contribution.

**Figure 1.**
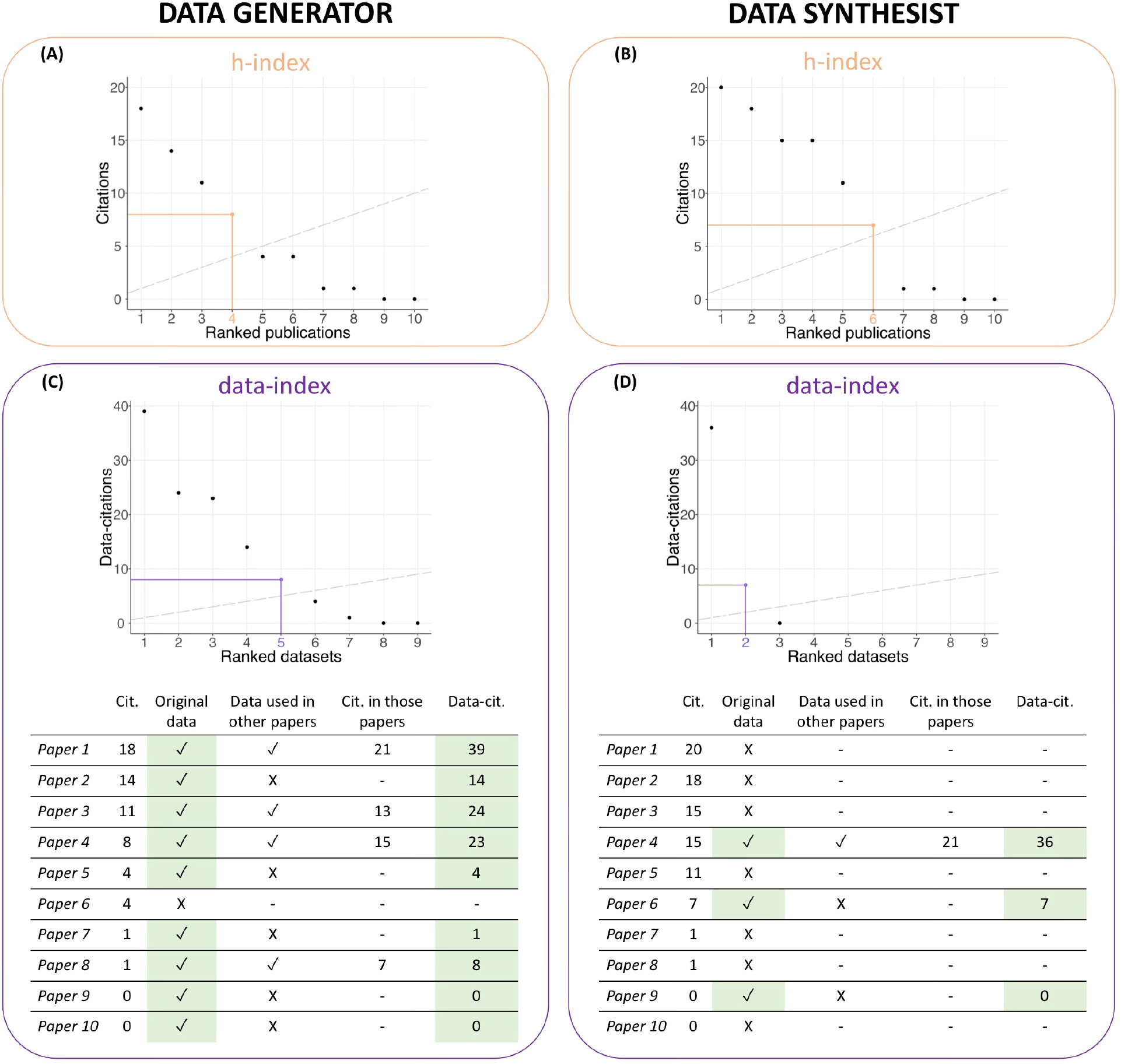
Scatterplots with publications ranked by citations to calculate h-index (A,B) and with datasets ranked by data-citations to calculate data-index (C,D). Dashed lines show identity lines and coloured lines show the final publication/dataset used to calculate the index value, which is also coloured. Tables show how to calculate the data-citations shown in the plots (C,D). Only highlighted rows, i.e. papers with original data, are included in the calculation. In this hypothetical example, the data generator has a lower h-index (4) (A) than the data synthesist (6) (B), but a higher data-index (6 vs 2) (C,D).

The data-index could easily be calculated automatically if authors were required to tag the DOIs of the publications that they extracted data from when they submitted their papers for review. It could be calculated retrospectively by generators and the authors that used the data tagging papers and validating each other’s claims, in a similar way to how authors currently claim their own publications online.

## Discussion

The data-index addresses two major issues in the fields of ecology and evolution: generating long-term datasets is undervalued and sharing data is disincentivised (Mills et al., 2015). Others have suggested changing authorship categories to better value data generators and address these issues (Ewers, Barlow, Banks-Leite, & Rahbek, 2019); the benefits of the data-index over changing authorship categories is that it is simple and can be implemented quickly and retrospectively.

Though our current system is satisfyingly simple, it is an oversimplification that hinders scientific progress and impact. Scientific contribution is diverse, and we need metrics that better value this diversity. The data-index is an example of a metric that can be used to complement metrics of publication impact and incentivise data sharing. More work should be done to generate indices for other aspects of scientific success, such as science communication (Sud & Thelwall, 2014), informing policy, mentoring scientists (Barres, 2013), or providing open access code, tools, or pipelines. Like many ecological and evolutionary processes, scientific success is multidimensional, and we must create a system that better values that.

## Conflict of Interest

None declared.

## Author Contribution

Amelia Hood: Conceptualization (lead); writing – original draft (lead); visualization (lead); methodology (lead); writing – review and editing (equal). William Sutherland: Writing – review and editing (equal); conceptualization (supporting); methodology (supporting).

## Notes

### Competing Interest Statement

The authors have declared no competing interest.

